# Using single-cell perturbation screens to decode the regulatory architecture of splicing factor programs

**DOI:** 10.1101/2025.02.07.637061

**Authors:** Miquel Anglada-Girotto, Samuel Miravet-Verde, Luis Serrano

## Abstract

Splicing factors shape the isoform pool of most transcribed genes, playing a critical role in cellular physiology. Their dysregulation is a hallmark of diseases like cancer, where aberrant splicing contributes to progression. While exon inclusion signatures accurately assess changes in splicing factor activity, systematically mapping disease-driver regulatory interactions at scale remains challenging. Perturb-seq, which combines CRISPR-based perturbations with single-cell RNA sequencing, enables high-throughput measurement of perturbed gene expression signatures but lacks exon-level resolution, limiting its application for splicing factor activity analysis. Here, we show that shallow artificial neural networks (ANNs) can estimate splicing factor activity from gene expression signatures, bypassing the need for exon-level data. As a case study, we map the genetic interactions regulating splicing factors during carcinogenesis, using the shift in splicing program activity –where oncogenic-like splicing factors become more active than tumor suppressor-like factors– as a molecular reporter of a Perturb-seq screen. Our analysis reveals a cross-regulatory loop among splicing factors, involving protein-protein and splicing-mediated interactions, with MYC linking cancer driver mutations to splicing regulation. This network recapitulates splicing factor modulation during development. Altogether, we establish a versatile framework for studying splicing factor regulation and demonstrate its utility for uncovering disease mechanisms.

**GRAPHICAL ABSTRACT.**
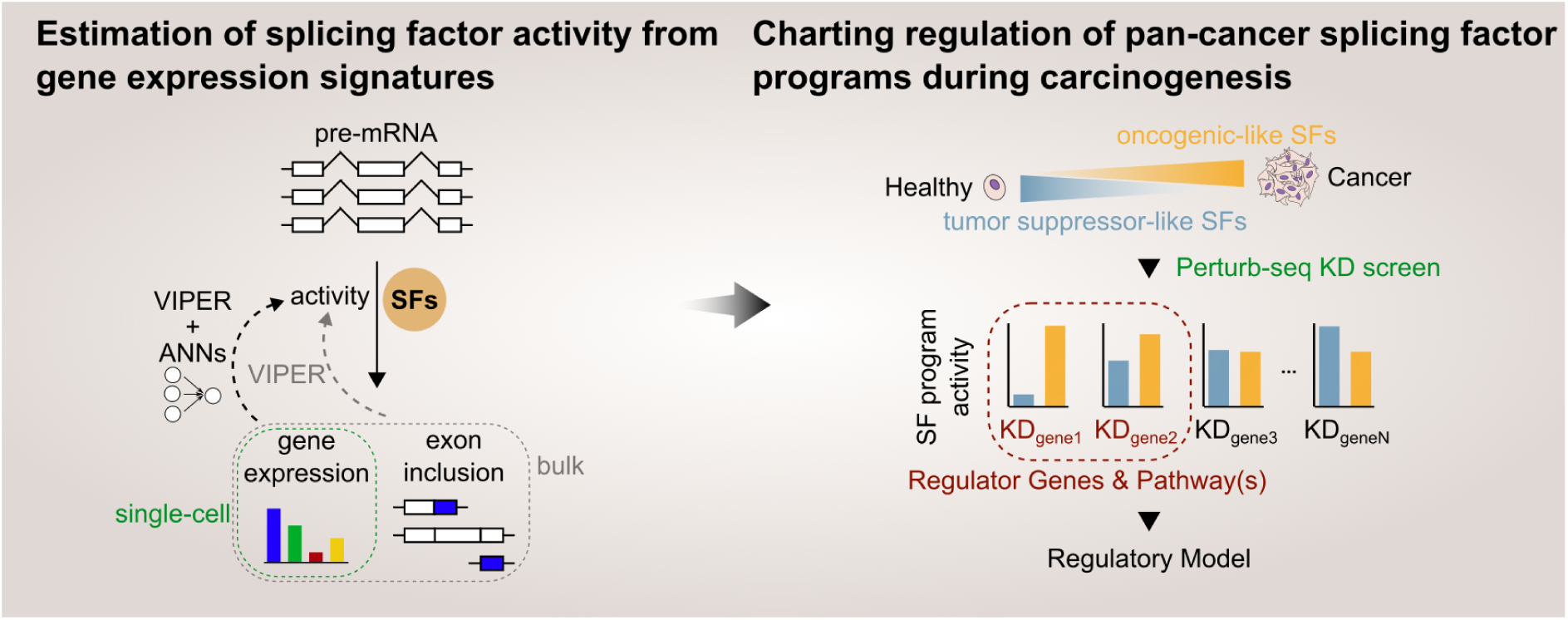

## INTRODUCTION

Splicing factors orchestrate the recognition and removal of introns in most pre-mRNAs, generating a pool of isoforms that contributes to maintaining homeostasis^1^. Numerous diseases, including cancer, co-opt splicing factors through mechanisms such as expression^2,3^, splicing^4^, post-translational modifications^5,6^, and protein-protein interactions^1^ to produce isoforms that drive disease progression^1^. Understanding how splicing factor activity becomes dysregulated is crucial for gaining insights into disease mechanisms and identifying potential therapeutic targets.

To measure how diverse molecular alterations influence splicing factors from a single omic dataset, we previously demonstrated that changes in exon inclusion provide a more accurate and comprehensive estimation of splicing factor activity than using their mRNA levels as a proxy because this approach captures multiple layers of regulation^7^. In this framework, “activity” refers to how much a given exon inclusion signature is enriched in the targets of a splicing factor, as quantified through virtual inference of protein activity by enriched regulon (VIPER^8,9^) (Fig. 1a). Thus, we hypothesize that integrating splicing factor activity analysis with transcriptomic outputs from large-scale genetic screens in disease-relevant experimental models could enable elucidating the regulatory interactions driving splicing factor dysregulation.

**Figure 1.**
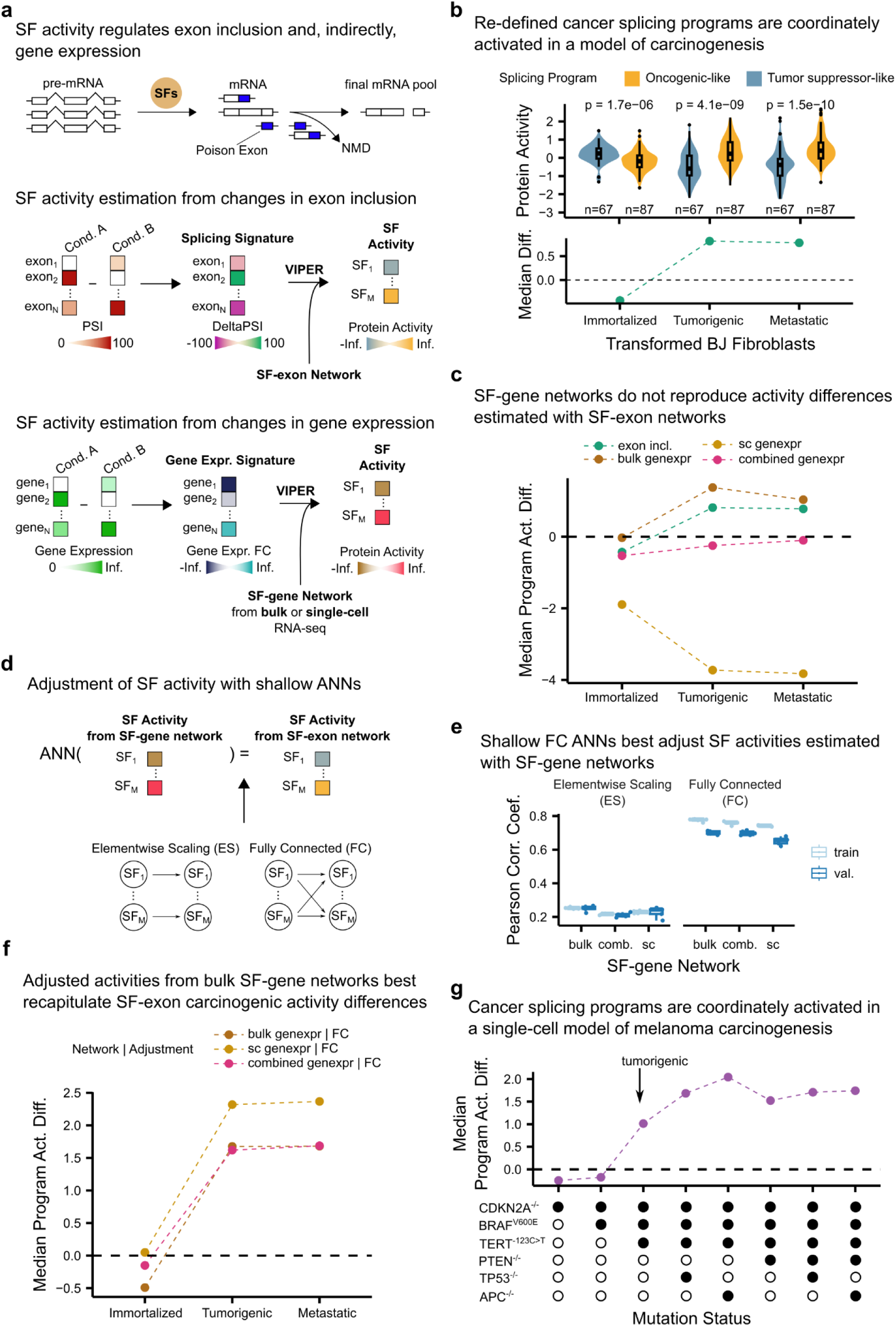
Quantifying splicing factor activity from gene expression data to use differential cancer splicing program activity as a reporter of carcinogenic transformation in single-cell transcriptomics datasets. **(a)** Splicing factor activity analysis with VIPER and splicing factor→exon networks. Splicing factors modulate exon inclusion, which can affect overall gene expression. For a signature of changes in exon inclusion, VIPER computes splicing factor activity given a network of interactions between splicing factors and exons. This approach can be adapted to estimate splicing factor activity from gene expression signatures. **(b)** Activity of cancer splicing programs across the carcinogenic stages of a fibroblast model^21^. Top, distributions of activities for each splicing factor with *p*-values from two-sided Wilcoxon rank sum tests. Bottom, median activity differences of oncogenic-like versus tumor suppressor-like splicing programs for each carcinogenic stage. **(c)** Median activity difference between cancer splicing programs across carcinogenic stages in Danielsson *et al.*’s model^21^. Activities were computed using different types of splicing factor networks: exon-based (“exon incl.”), bulk RNA sequencing gene-based (“bulk genexpr”), single-cell RNA sequencing gene-based (“sc genexpr”), bulk and single-cell RNA sequencing gene-based (“combined genexpr”). We consider exon-based splicing factor networks as our ground truth, hence, the closer that program activity differences are to “exon incl.” lines the better. **(d)** Splicing factor activities computed using gene-based splicing factor networks can be adjusted with shallow artificial neural networks (ANN) to resemble those computed using exon-based splicing factor networks, which we consider ground truth. **(e)** Pearson correlation coefficients between exon-based and gene-based splicing factor activities adjusted with shallow ANNs trained using CCLE data. Element-wise scaling ANNs consist of a single layer of parameters that scale each gene-level activity independently while fully connected ANNs consider gene-level activities for all splicing factors to adjust them. **(f)** Median activity difference between cancer splicing programs adjusted with fully connected ANNs across the carcinogenic stages of Danielsson *et al.*’s model^21^. Activities were computed considering different gene-based splicing factor networks. **(g)** Median activity difference between cancer splicing programs computed using bulk gene-level networks adjusted with fully connected ANNs. Activities were computed across the carcinogenic stages of Hodis *et al.^24^*’s model, which transformed melanocytes into melanoma through iterative mutations (x-axis). Arrow, the stage at which transformed cells acquire tumorigenic capacity. In box and whisker plots in panels (b) and (e) the median is marked by a horizontal line, with the first and third quartiles as box edges. Whiskers extend up to 1.5 times the interquartile range, and individual outliers are plotted beyond.

**Supplementary Figure 1.**
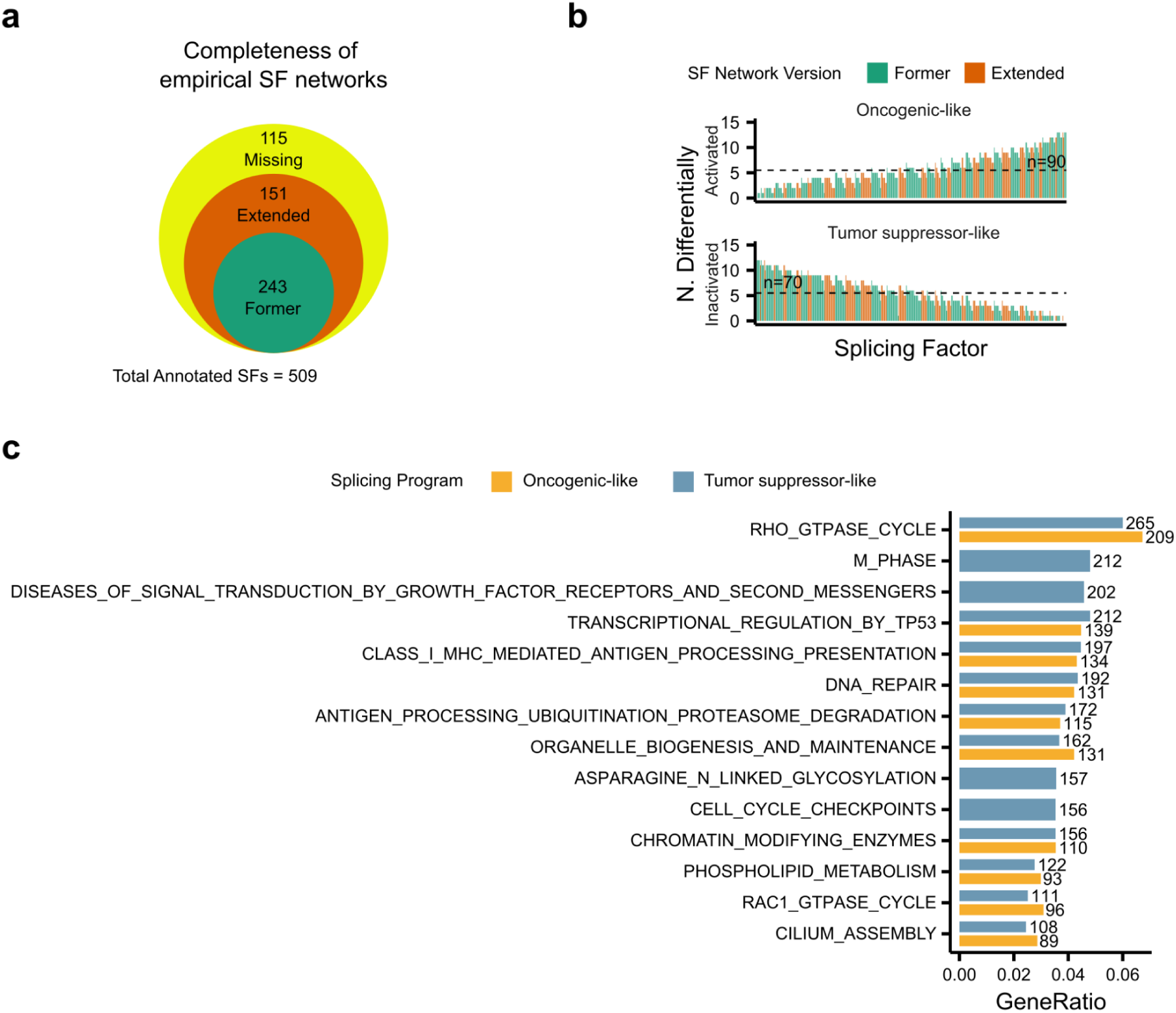
Benchmarking extended splicing factor networks and re-defining cancer splicing programs. **(a)** Venn diagram of the number of splicing factors considered in the former and extended versions of empirical splicing factor networks compared to the total of annotated splicing factors. **(b)** Re-definition of cancer splicing programs with extended splicing factor networks. Number of cancer cohorts in which a splicing factor is recurrently activated (top) or inactivated (bottom). Cancer splicing programs are defined with those splicing factors recurrently active in at least 5 cancer types. **(c)** Overrepresentation analysis (ORA) of genes corresponding to target exons of splicing factors in re-defined cancer splicing programs. Gene sets from ReactomeDB (n=1,604). We only show the top 10 significantly enriched sets (FDR < 0.05, ORA) sorted by gene ratio combining the enriched gene sets for both splicing programs. A total of 98 and 262 gene sets were enriched for oncogenic-like and tumor suppressor-like splicing factors. Those gene sets missing one cancer splicing program imply that the given gene set was not significantly enriched in the missing program.

While genetic screens with exon inclusion resolution are limited and often heterogeneous, the availability of large-scale perturbation screens with single-cell transcriptomics outputs, such as Perturb-seq, is rapidly expanding^10–12^. These assays provide the throughput and causal connections necessary for systematically investigating splicing factor regulation^13^. However, published Perturb-seq screens use 10X droplet-based methods with 3’-end transcript capture, providing data with limited insight into the alternative splicing changes essential for the current implementation of splicing factor activity analysis^14^.

In cancer, emerging evidence indicates that groups of splicing factors with oncogenic-like or tumor suppressor-like behavior change in a coordinated manner across multiple cancer cohorts following carcinogenic transformation^3,7,15,16^. In particular, several studies have identified MYC, a widely-studied oncogenic transcription factor, as an important regulator of these splicing factor programs^3,17–19^. However, beyond MYC, the involvement of other genes and pathways that may cooperate in regulating these splicing programs has remained largely unexplored due to limitations in studying carcinogenic splicing factor regulation at scale.

Here, we show that single-layer artificial neural networks (ANNs) in combination with Perturb-seq studies enable the use of single-cell transcriptomics to infer splicing factor activity in carcinogenesis. We validated this approach by identifying the coordinated change in activity of splicing factors with oncogenic-like and tumor suppressor-like behavior across multiple cancer cohorts following carcinogenic transformation, previously found using inclusion/exclusion of exons^7^. Our analysis uncovers a cross-regulatory loop among splicing factors involving protein-protein and splicing-mediated interactions and identifies MYC as the key pathway linking cancer driver mutations to splicing factor regulation. This regulatory architecture mirrors the developmental modulation of splicing factors, suggesting that cancer hijacks this pre-existing regulatory program. Our approach establishes a framework for investigating splicing factor regulation through single-cell transcriptomics, offering a versatile tool for studying cellular processes and splicing-associated diseases.

## RESULTS

### Estimation of splicing factor activity from gene expression signatures using shallow artificial neural networks

To develop a methodology for studying splicing factor activity from single-cell data without relying on exon inclusion/exclusion profiles, we first require a reliable reference dataset for validation. Previously, we showed that by analyzing splicing factor→exon networks derived from bulk RNA sequencing datasets we can link changes in exon inclusion to splicing factor activity, capturing multiple layers of regulation of splicing factors^7^. To expand the coverage of splicing factor activity estimation to be used to compare with that inferred from single-cell data, we processed the recent dataset from Rogalska *et al.*^20^, where HeLa cells were transcriptomically profiled after silencing splicing factors (see Methods). This integration adds 151 splicing factors to our analysis (n_total_=394), leaving 115 out of 509 annotated human splicing factors without a network of targets (Supplementary Fig. 1a).

By comparing splicing factor activities in primary tumors and their solid tissue normal counterparts across 14 cancer cohorts (FDR < 0.05, two-sided Wilcoxon rank sum test; see Methods), we identified 90 splicing factors as recurrently active (oncogenic-like) and 70 as recurrently inactive (tumor suppressor-like) in at least 5 cancer cohorts (Supplementary Fig. 1b; Supplementary Table 1; see Methods). These cancer splicing programs are enriched in proliferation-related targets (FDR < 0.05, Overrepresentation Analysis (ORA)), supporting their influence on the proliferative capacity of cancer cells (Supplementary Fig. 1d). As previously observed^7^, analyzing the data from Danielsson *et al.* profiling transformed BJ fibroblasts as they progressed through immortalized, tumorigenic and metastatic stages^21^ revealed that oncogenic-like splicing factors become more active than tumor suppressor-like ones during the transition from immortalized to tumorigenic stages (Fig. 1b). The median activities of oncogenic-like and tumor suppressor-like programs could serve as a molecular reporter of carcinogenic regulation, with negative scores indicating greater activity of the tumor suppressor-like program (pre-tumorigenic stage), while positive scores reflecting the dominance of the oncogenic-like program (tumorigenic stage) (Fig. 1b).

To study splicing factor regulation through splicing factor activity analysis in datasets where exon inclusion cannot be quantified, we considered that splicing factors indirectly regulate gene expression. The inclusion or exclusion of certain exon combinations affects mRNA stability and can trigger nonsense-mediated decay (NMD), affecting the final pool of mRNAs and measured gene expression^22,23^. Additionally, all those exon inclusion changes that do not trigger NMD may also alter the activity of downstream genes that could influence transcription and or degradation of mRNA. Hence, splicing factor activities could potentially be estimated from the changes in gene expression between conditions, interpreted with VIPER and networks linking splicing factors to genes (Fig. 1a).

We evaluated splicing factor→gene networks generated from datasets measuring transcriptomic changes when perturbing splicing factors using bulk or single-cell RNA sequencing. To construct splicing factor→gene networks from bulk RNA sequencing data, we leveraged the perturbation datasets originally used to generate splicing factor→exon networks. Here, genes were defined as targets if their expression changed by at least |1| log2(TPM+1) upon perturbation of a splicing factor (see Methods). In this setting, gene expression signatures replaced exon inclusion data to estimate splicing factor activity. Additionally, we generated splicing factor→gene networks using Perturb-seq datasets from Replogle *et al.*^10^, which involved single-cell RNA sequencing of two cell lines, K562 and RPE1, following the perturbation of all expressed (n=9,867) and all essential genes (n=2,285), respectively (see Methods). To maximize the coverage, we combined the bulk and single-cell networks into a third type of splicing factor network, considering as many splicing factor→gene interactions as possible (n_unique interactions_: bulk = 977,505; single-cell = 321,607; combined = 1,248,595) (see Methods).

For each gene-based network (i.e., bulk, single-cell, and combined), we evaluated how well estimated splicing factor activities matched the median activity differences between cancer splicing programs during carcinogenesis derived from exon-based networks, as defined above. We found that none of the gene-based networks fully recapitulated the activities estimated using exon-based networks (Fig. 1c). Of the three datasets, bulk RNA sequencing gene-based networks produced activity differences most closely aligned with exon-based networks (Euclidean distance = 0.73). However, in the immortalized stage, the median program activity difference estimated from bulk gene-based networks was near zero, whereas exon-based networks –our ground truth– showed a negative activity difference (Fig. 1c). Thus although splicing factors influence gene expression, gene-based networks cannot fully capture cancer splicing program activity differences.

Nevertheless, the relatively small distance (Euclidean distance = 0.73) with exon-based activities of bulk gene-based networks suggests that gene-based activities can be mathematically adjusted to better match exon-based ones. Artificial neural networks (ANNs) offer a powerful solution for modeling such complex relationships by processing input data through weighted connections to predict outputs. We implemented a one-layer ANN with a fully connected layer, which considers the interactions between all estimated splicing factor activities and, as a baseline, an element-wise multiplication layer, where each splicing factor activity is scaled by a weight, equivalently to fitting a linear model for each factor. We trained these shallow ANNs using splicing factor activities estimated from 1,019 cancer cell lines in the Cancer Cell Line Encyclopedia (k-fold cross-validation = 5). The input data consisted of splicing factor activities derived from either bulk, single-cell or combined gene-based splicing factor networks. The output data comprised splicing factor activities estimated using exon-based networks (Fig. 1d; see Methods). Adjusting activities using the element-wise layer resulted in underfitting the data, with an overall training and validation Pearson Corr. Coef.≃0.2. This suggests that the ANN architecture requires a more complex architecture to capture this relationship. Indeed, adjusting activities with a fully connected layer achieved high performance with minimal overfitting for all gene-based networks (overall Pearson Corr. Coef. training≃0.75, and validation≃0.65) (Fig. 1e).

In the context of carcinogenesis as defined above, splicing factor activities estimated using bulk gene-based networks and adjusted with pre-trained fully connected ANNs closely mirrored the behavior of exon-based networks. Specifically, the median program activity differences were negative in the immortalized stage and remained positive in the tumorigenic and metastatic stages (Fig. 1f). This demonstrates that adjusted activities derived from gene expression signatures are accurate enough to use program activity differences as reliable reporters of carcinogenic regulation. In principle, this extends splicing factor analysis to datasets where exon inclusion quantification is not feasible.

### Validating cancer splicing program activity difference as a reporter of carcinogenic transformation in single-cell data

To validate the use of cancer splicing program activity difference as a reliable reporter of carcinogenic transformation in single-cell data, we analyzed an experimental model of melanoma carcinogenesis^24^. In Hodis *et al.^24^*’s study, primary melanocytes were genomically engineered in iterative steps to acquire invasive melanoma phenotypes. Each combination of mutations was characterized through single-cell RNA sequencing.

Using our ANN-based approach, we estimated splicing factor activities for each carcinogenic stage and computed the median activity difference between splicing factors in the oncogenic-like and tumor suppressor-like splicing programs. The median activity of oncogenic-like splicing factors surpassed that of tumor suppressor-like splicing factors only after the introduction of CDK2A^-/-^, BRAF^V600E^, and TERT^-123C>T^ mutations, which confer tumorigenic capacity (Fig. 1g). This switch in activity mirrors the pattern observed in the bulk RNA sequencing-based carcinogenesis experiment, demonstrating the robustness and generalizability of our approach. These results confirm that our pipeline for estimating splicing factor activities from gene expression signatures provides a reliable framework for studying the carcinogenic regulation of cancer splicing programs in both bulk and single-cell transcriptomic datasets.

### Protein-protein and splicing-mediated cross-regulation among splicing factors coordinately activates oncogenic-like splicing factors and inactivates tumor suppressor-like factors

Studying the regulation of cancer splicing programs during carcinogenesis requires experimentally assessing how genetic perturbations influence their activity, ideally within a cellular context representing a pre-tumorigenic stage. Replogle *et al.* conducted a Perturb-seq screen that measured gene expression changes resulting from CRISPR silencing of 2,285 genes in RPE1 cells, an immortalized cell line^10^. Using our gene expression-based pipeline, we quantified the extent to which each knockdown alters the activity of cancer splicing programs. This approach offers insights into the regulatory mechanisms underlying these splicing programs. In particular, this analysis allows identifying gene perturbations that induce a switch in splicing program activity by coordinately activating oncogenic-like splicing factors while inactivating tumor suppressor-like factors, mirroring the transition observed in carcinogenesis experiments (Fig. 2a).

**Figure 2.**
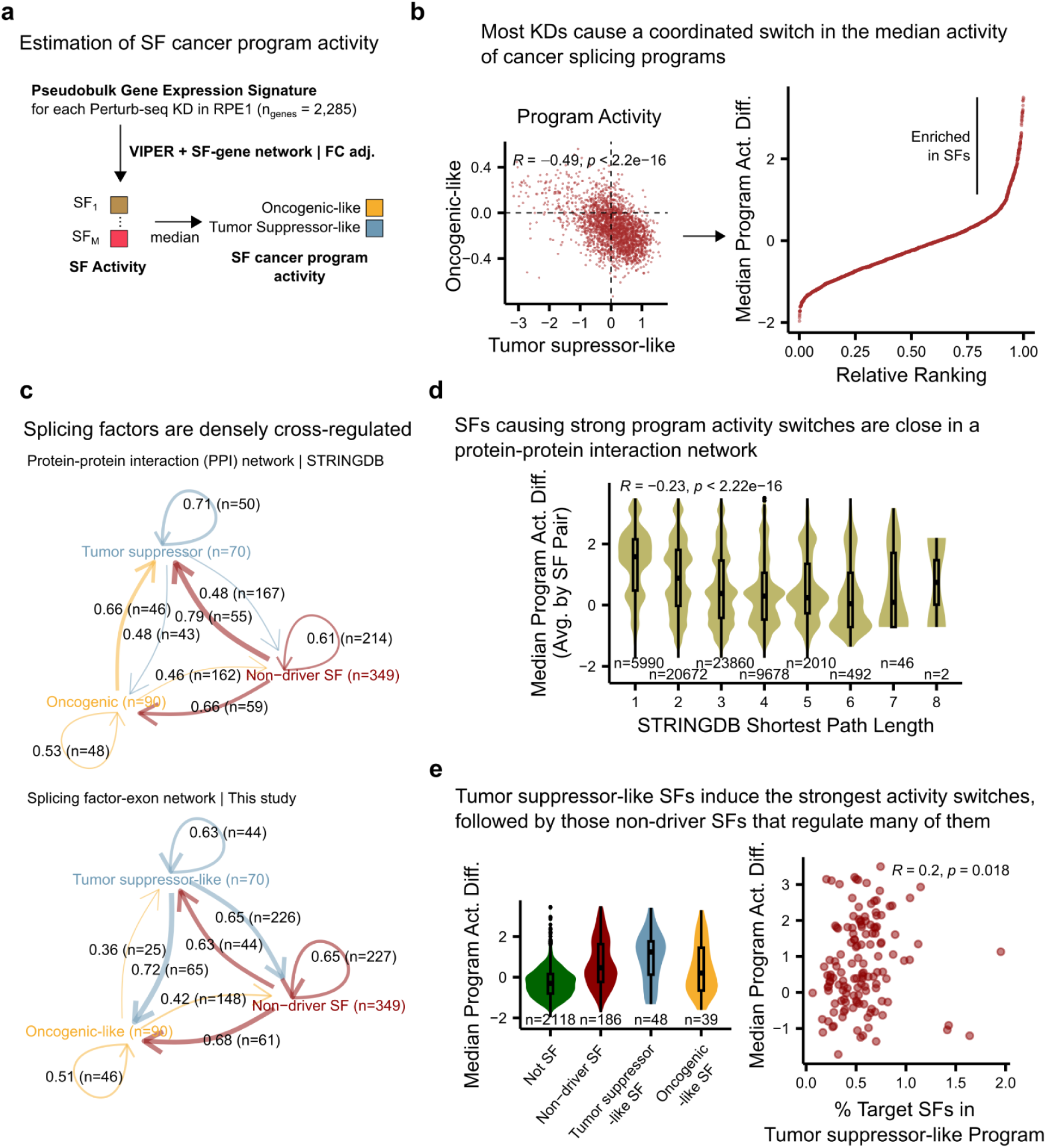
Systematic identification of cancer splicing program regulators through splicing factor activity analysis of a Perturb-seq dataset. **(a)** Workflow to estimate median activity differences between cancer splicing programs for each knockdown in Replogle *et al.*’s Perturb-seq dataset using the immortalized RPE1 cell model^10^. **(b)** Distributions of median activities for each cancer splicing program (left) and their difference (right). Top-left, Pearson correlation coefficient, and corresponding p-value. **(c)** Splicing factor cross-regulation through protein-protein and splicing factor→exon interactions. Each arrow indicates the ratio and number of splicing factors of the target class regulated by the source class. Arrow thickness, splicing factor target ratio. **(d)** Distributions of median program activity differences between cancer splicing programs of pairs of splicing factors versus their shortest path distance in the STRINGDB protein-protein interaction network. Top, Spearman correlation coefficient, and corresponding p-value. **(e)** Left, distributions of median program activity differences between cancer splicing programs for each splicing factor class and the rest of the genes. Right, median program activity differences between cancer splicing programs for knockdowns of “non-driver-like” splicing factors versus the percentage of target exons of each splicing factor corresponding to tumor suppressor-like splicing factors. In box and whisker plots in panels (d) and (e) the median is marked by a horizontal line, with the first and third quartiles as box edges. Whiskers extend up to 1.5 times the interquartile range, and individual outliers are plotted beyond.

**Supplementary Figure 2.**
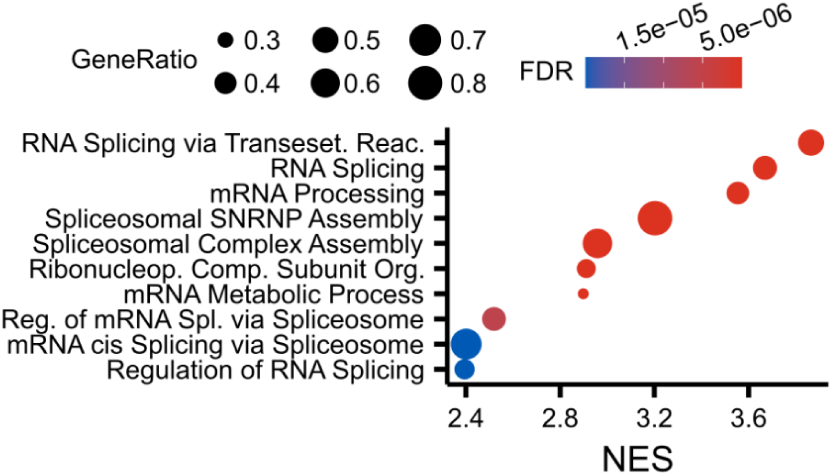
Enrichments of activity differences between cancer splicing programs. Top 10 significantly enriched (FDR < 0.05, GSEA) GO biological processes based on Perturb-seq knocked-down genes sorted by corresponding median activity difference between cancer splicing programs.

To characterize the perturbed genes capable of regulating cancer splicing programs, we performed gene set enrichment analysis (GSEA), sorting the genes by the median program activity differences between oncogenic-like and tumor suppressor-like splicing factors resulting from each gene silencing in the Perturb-seq screen. Most enriched terms (FDR < 0.05, GSEA) were related to splicing, indicating that splicing factors themselves are important in regulating the carcinogenic activation of cancer splicing programs (Supplementary Fig. 2). This suggests that cancer leverages splicing factor cross-regulation to collectively control their activity.

Next, we investigated whether functional interactions among splicing programs could explain the coordinated regulation of oncogenic-like and tumor suppressor-like splicing programs observed during carcinogenesis. If such cross-regulation exists, perturbations that alter the activity of one program would inversely affect the activity of the other. Conversely, in the absence of cross-regulation, perturbations affecting one program would show no correlation with the other. Analyzing the median splicing factor activities of each program for all knockdowns revealed a strong inverse relationship: most perturbations activating the oncogenic-like program simultaneously inactivate the tumor suppressor-like program, or vice-versa (Pearson Corr. Coef. = −0.49, p < 2.2·10^-16^) (Fig. 2b). This functional entanglement between cancer splicing programs suggests that the activity switch observed during carcinogenesis is driven by cross-regulatory interactions among splicing factors.

Splicing factors regulate their activity through interactions at multiple levels, including protein-protein interactions (PPIs) and splicing factor→exon interactions. To evaluate the extent to which these cross-regulatory interactions could control cancer splicing programs, we analyzed the proportion of splicing factors in each class –oncogenic-like, tumor suppressor-like, and non-driver-like– regulated by another splicing factor class. This analysis was performed using the STRINGDB PPI network^25^ and the splicing factor→exon networks generated in this study. Both types of networks are highly interconnected with approximately 50% of splicing factors in each class regulated by factors from other classes (Fig. 2c). For example, at the PPI level, tumor suppressor-like splicing factors interact with 48% of oncogenic-like splicing factors, while oncogenic-like splicing factors interact with 66% of tumor suppressor-like factors. Similarly, at the splicing level, non-driver-like splicing factors regulate more than half of the splicing factors involved in cancer splicing programs (Fig. 2c). These findings highlight the extensive cross-regulation among splicing factors across both protein-protein and splicing-mediated interactions, suggesting a tightly interconnected regulatory network that governs cancer splicing programs.

To study the influence of PPI networks on the regulation of cancer splicing programs, we analyzed the shortest path length distributions between splicing factor pairs in the STRINGDB network that were also perturbed in the Perturb-seq dataset (see Methods). We then correlated these distances with the average median program activity differences derived from the Perturb-seq assay. These results revealed that activity differences decrease as the distance between each splicing factor pair increases (Spearman Corr. Coef. = −0.23, p < 2.2·10^-16^), plateauing at 0 when the splicing factor pair is three or more steps apart (Fig. 2d). This suggests direct PPIs among splicing factors play an important role in the coordinated regulation of cancer splicing programs.

Next, we examined how splicing factor→exon interactions influence cancer splicing program regulation. We find that oncogenic-like splicing factors did not result in strong median program activity differences, consistent with the inactivating fashion of the knockdown screen analyzed. In contrast, knocking down tumor suppressor-like splicing factors caused the largest program activity switches (median program activity difference = 1.24) compared to the rest of the splicing factor classes and genes (Fig. 2e). However, some non-driver-like splicing factors exhibited program activity differences comparable to tumor suppressor-like splicing factors, despite not displaying recurrent activation or inactivation across cancer cohorts (Supplementary Table 1). We hypothesized that these non-driver-like factors may indirectly regulate program switches by controlling exon inclusion in tumor suppressor-like splicing factors. Supporting this, the greater the fraction of tumor suppressor-like splicing factors regulated by a non-driver-like factor, the stronger the program activity difference observed upon its silencing (Spearman Corr. Coef = 0.2, p = 0.018) (Fig. 2e, right).

These results suggest that cancer splicing programs operate within a cross-regulatory feedback loop driven by both protein-protein and splicing-mediated interactions. This loop facilitates the coordinated activation of oncogenic-like splicing factors and the inactivation of tumor suppressor-like splicing factors. This dense and redundant cross-regulation among splicing factors provides multiple points of control, enabling cancer cells to effectively and adaptively exploit the splicing machinery.

### Systematic re-discovery of MYC as a regulator of cancer splicing programs

While numerous splicing factors can induce the coordinated regulation of cancer splicing programs, most cancer driver genes listed in the COSMIC Cancer Gene Census^26^ are not splicing factors (48 of 733 are splicing factors). In carcinogenesis experiments, we observe a switch in splicing program activity even though the cancer driver alterations introduced do not involve splicing factors directly. This suggests that specific pathways mediate the translation of oncogenic lesions into changes in cancer splicing programs. To systematically identify these mediator pathways, we analyzed pathway normalized enrichment scores (NESs) during carcinogenesis and following gene perturbations (Fig. 3a).

**Figure 3.**
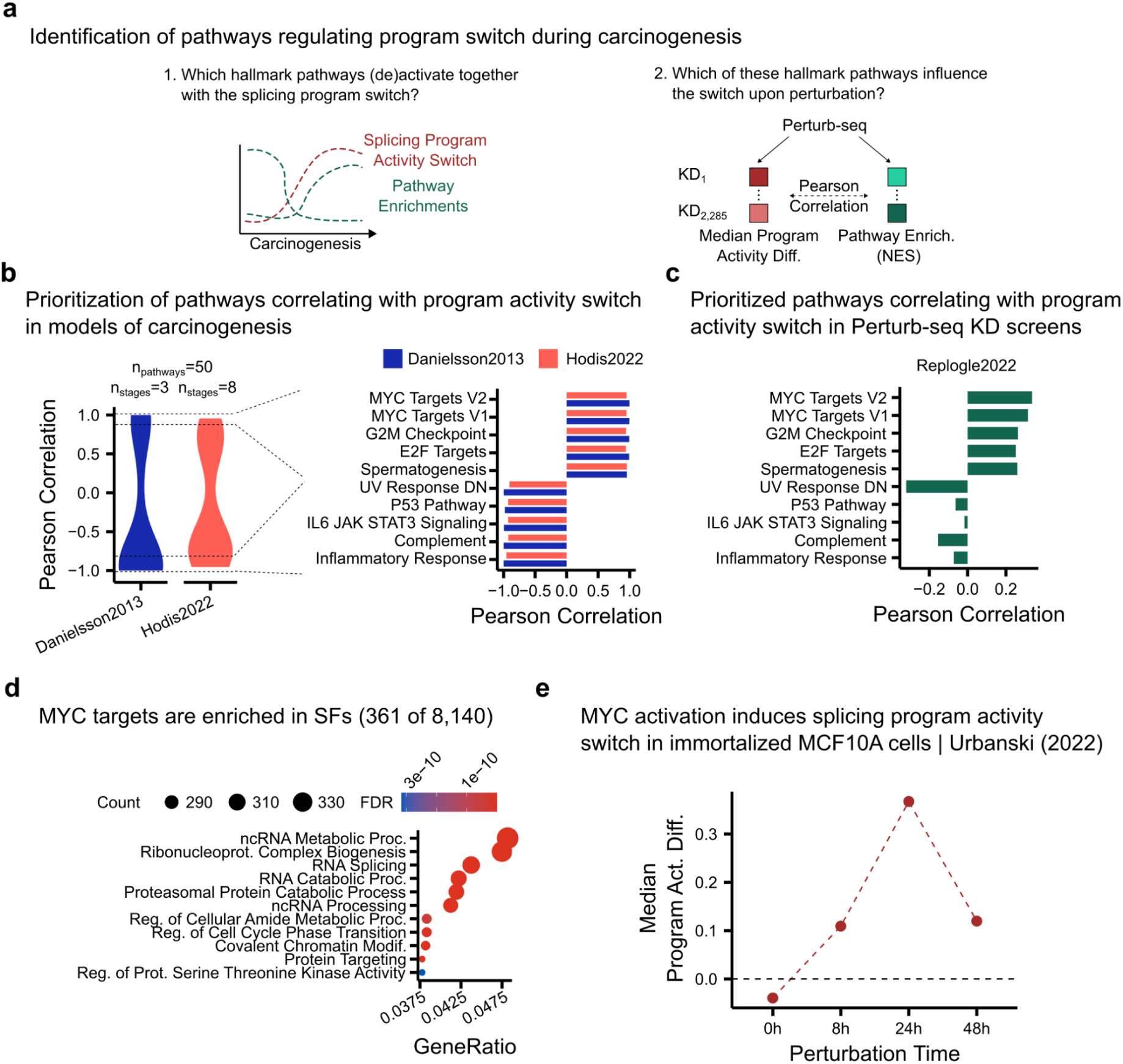
Identification and validation of MYC as the regulator translating carcinogenic alterations into cancer splicing program regulation. **(a)** Prioritization of pathways with the potential to translate cancer driver alterations into regulation of cancer splicing programs. **(b)** Distribution (left) and top 10 (right) MSigDB Hallmark pathways whose GSEA NES correlates with median activity differences between cancer splicing programs across carcinogenic stages of models from Danielsson *et al.* and Hodis *et al*. **(c)** Pearson correlations between NES of pathways prioritized in (b) and median activity differences between cancer splicing programs for each knockdown in Replogle *et al.* Perturb-seq dataset using the immortalized RPE1 cell model. **(d)** Top 10 enriched (FDR < 0.05, ORA) gene sets sorted by gene ratio for MYC target genes listed in the CHEA Transcription Factors database^28^. **(e)** Normalized median activity differences between cancer splicing programs in immortalized MCF10A cells carrying a MYC activation system inducible with 4-hydroxytamoxifen (4-OHT). Estimated splicing factor activities were normalized by subtracting the activities of the negative control (MCF10 cells without the inducible system treated with 4-OHT at matching time points).

**Supplementary Figure 3.**
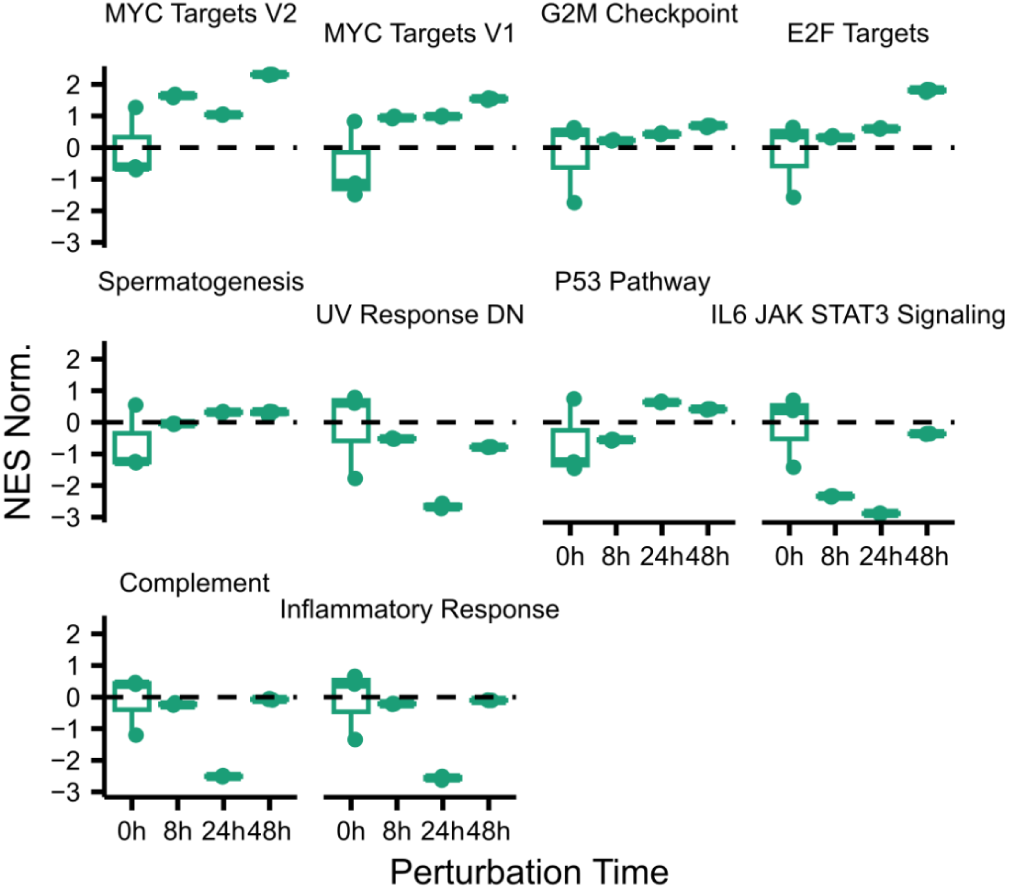
GSEA of changes in expression upon induced MYC activation. Normalized NES from GSEAs for each pathway prioritized in Fig. 3b in immortalized MCF10A cells inducibly activating MYC upon 4-OHT treatment. We normalized NES by subtracting NES from the control condition, MCF10A cells without the inducible system treated with 4-OHT for the same time points. In box and whisker plots, the median is marked by a horizontal line, with the first and third quartiles as box edges. Whiskers extend up to 1.5 times the interquartile range, and individual outliers are plotted beyond.

First, we reasoned that potential regulatory pathways of cancer splicing programs should become active or inactive concomitantly to the switch in the splicing programs throughout carcinogenesis. We obtained NESs for the pathways in the MSigDB hallmark gene sets^27^ for both of the carcinogenesis experiments analyzed and correlated these NES with median cancer splicing program activity differences previously computed (see Methods). From the distributions of correlations, we shortlisted the top and bottom five pathways with the strongest concordant correlations in both experiments as candidate regulatory pathways (Fig. 3b).

To distinguish which of the shortlisted pathways regulates cancer splicing programs, we should perturb one or more genes in every candidate pathway and measure how this influences the splicing program activity switch. Perturb-seq screens measure the changes in gene expression upon silencing many different genes. Each perturbation may alter different pathways directly and indirectly. Hence, correlating pathway NES with median program activity differences across a large number of KDs should indicate which shortlisted pathways are more likely to regulate cancer splicing programs. Pathways “MYC Targets V2” and “MYC Targets V1” exhibit the strongest correlations among shortlisted pathways, suggesting MYC as the mediator regulator (Fig. 3c). In line with this, 361 out of 8,140 chromatin immunoprecipitation-based MYC targets are splicing factors^28^. Considering there are 509 annotated splicing factors, this represents a significant enrichment (FDR < 0.05, ORA) (Fig. 3d; see Methods). These results highlight MYC as the potential regulator connecting cancer driver alterations to cancer splicing programs.

Although MYC is known to regulate cancer splicing programs, we sought to validate its role specifically within the context of splicing factor activity analysis. To this end, we analyzed transcriptomic data from Urbanski *et al.*, who characterized immortalized MCF10A cells with an inducible MYC activation system^3^. Consistent with patterns observed in carcinogenesis, MYC induction resulted in a positive normalized program activity difference, reflecting the activation of oncogenic-like splicing factors and inactivation of tumor suppressor-like splicing factors (Fig. 3e; see Methods). As a control for our systematic prioritization method, we also confirmed that MYC induction increased NESs for MYC-related pathways (Supplementary Fig. 3). These findings demonstrate that MYC regulation is sufficient to drive the coordinated activation and inactivation of cancer splicing programs during carcinogenesis.

These results establish MYC as a central regulator connecting cancer driver alterations to cancer splicing programs during carcinogenesis.

### Carcinogenic regulation of cancer splicing programs mirrors developmental splicing control

Oncogenic processes repurpose existing cellular programs to promote cancer hallmarks, including embryonic programs of dedifferentiation^29^. We then wondered whether the differential activation of cancer splicing programs observed during carcinogenesis reflects the hijacking of pre-existing developmental splicing programs.

Cardoso-Moreira *et al.* analyzed the transcriptomes of nine human tissues across developmental stages, from embryonic to adult^30^. We processed these data to estimate the activities of cancer splicing programs (see Methods). To explore the relationship between cancer splicing program activation and developmental timing, we grouped samples from each tissue into 10 bins based on days post-conception. Unlike in carcinogenesis, we observed that tumor suppressor-like splicing factors exhibit higher activity than oncogenic-like splicing factors during developmental differentiation (Fig. 4a). This is reflected in the negative correlation between cancer splicing program activity differences and developmental progression across most tissues (overall Spearman Corr. Coef. = −0.42, p = 9.7·10^-5^) (Fig. 4a). This suggests that cancer splicing programs originate from embryonic splicing programs active during early development.

**Figure 4.**
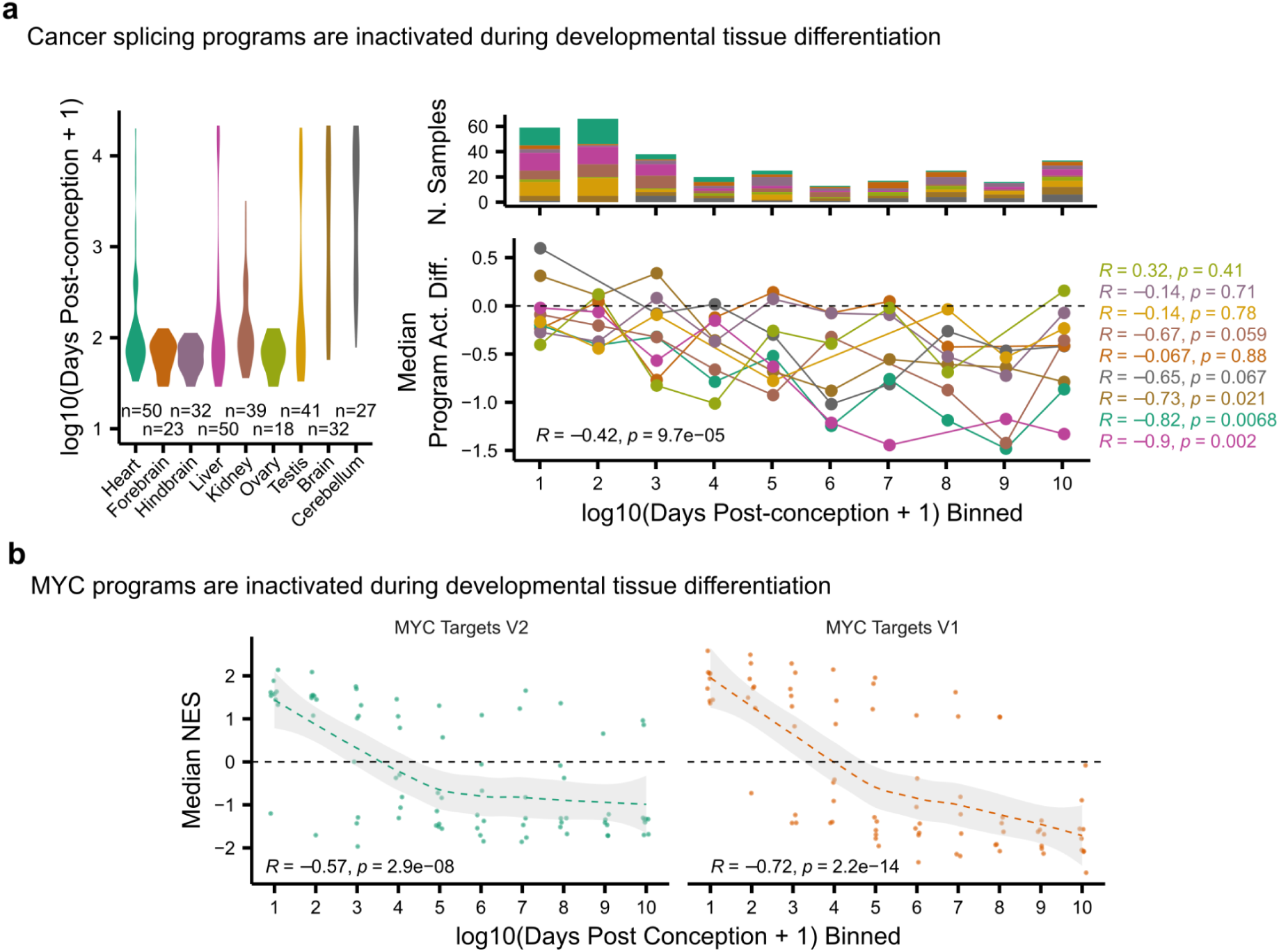
The activity of cancer splicing programs during development. **(a)** Splicing factor activity analysis of developmental samples from different stages of tissue differentiation. Left, distributions of developmental time (days post conception) for the samples of each tissue. Right-top, number of samples from each tissue at each developmental time bin. Right-bottom, median activity differences between cancer splicing programs for each tissue across developmental time bins. Labels, Spearman correlation coefficients for all (plot bottom) and each tissue (right), and corresponding p-values are reported. **(b)** Median NES from GSEAs for MYC target pathways across binned developmental time for the 9 tissues sampled in Cardoso-Moreira *et al.* The samples and time bins are the same ones used in panel (a). Bottom labels, Spearman correlation, and corresponding p-values. Dashed line represents the fitted trend line using the LOESS method, and the shaded region depicts the 95% confidence interval, as determined by *ggplot2::geom_smooth(method = ‘loess’)* R function.

MYC, a transcription factor involved in cellular growth and proliferation, is a well-established regulator of pluripotency in healthy contexts^31^ and is frequently upregulated in cancer^32^. This suggests cancer cells hijack MYC’s regulatory network to control cancer splicing programs. Supporting this hypothesis, MYC-target pathway enrichments decrease during healthy tissue differentiation, consistent with the reduced activity of oncogenic-like splicing factors compared to tumor suppressor-like factors during development (Fig. 4a). Specifically, NESs for MYC-target pathways (V1 and V2) showed negative correlations with developmental time bins (Spearman Corr. Coef. = −0.57 and −0.72) (Fig. 4b).

These findings further support MYC as a central regulator of cancer splicing programs and suggest that these programs are derived from pre-existing developmental regulatory programs.

## DISCUSSION

Splicing factors regulate the isoform pool expressed by most genes across all cellular contexts. Many diseases exploit this fundamental function to drive hallmark processes. Understanding the regulation of splicing factors is therefore critical to improving diagnostics and treatments. While aberrant splicing factor activity can be accurately assessed from exon inclusion signatures, only single-cell transcriptomics provides the necessary throughput to systematically map splicing factor regulation through large-scale genetic screens. However, the limited transcript coverage of single-cell assays restricts the analysis to gene-level signatures, making traditional exon-based splicing factor activity analysis infeasible and leaving valuable insights untapped.

In this study, we developed an approach to perform splicing factor activity analysis using gene expression signatures, focusing on understanding the carcinogenic regulation of cancer splicing programs. By extending splicing factor→exon networks with recent perturbation datasets, we enabled activity estimation for 151 additional splicing factors, allowing us to refine cancer splicing programs. To address the limitations of exon inclusion data in single-cell transcriptomics, we used shallow ANNs to estimate splicing factor activities from gene expression changes. Although more complex ANNs could have been applied, this simplified approach effectively validated the coordinated activation of oncogenic-like splicing factors and inactivation of tumor suppressor-like factors as a reliable molecular reporter of carcinogenic transformation in both bulk and single-cell datasets. Using this reporter, we analyzed Perturb-seq experiments in a pre-tumorigenic cellular model to identify potential regulators of the switch in cancer splicing program activity observed during carcinogenesis. Further, we found that cancer splicing programs exhibit inverse regulation during developmental tissue differentiation, suggesting that cancer hijacks pre-existing embryonic pathways to drive disease progression.

Our analyses revealed a dense and redundant cross-regulation between splicing factors, forming a cross-regulatory loop that underpins the coordinated regulation of cancer splicing programs. We identified protein-protein and splicing-mediated interactions as relevant layers supporting this cross-regulation. However, additional molecular mechanisms –such as post-transcriptional and translational modifications^5,6,33,34^– may also contribute but require further investigation. This extensive cross-regulation enables multiple splicing factors to trigger the switch in program activity, representing a potential therapeutic target. Notably, carcinogenesis models showed this activity switch despite not having perturbed splicing factors directly. Thus, we investigated upstream pathways connecting cancer driver lesions to splicing factor regulation. Through analyses of carcinogenesis and Perturb-seq datasets, MYC emerged as the strongest candidate regulator. Supporting this, experimental activation of MYC induced the splicing program activity switch. These findings position MYC as a master regulator that connects oncogenic lesions to splicing factor networks, driving the transition to a tumorigenic state^3,19^.

Considering these results, we propose the following model of cancer splicing program regulation during carcinogenesis (Fig. 5). In healthy cells, diverse oncogenic lesions activate the MYC pathway, altering the expression of key “initiator” splicing factors. This rewires the splicing factor cross-regulatory network, leading to the coordinated activation of oncogenic-like and inactivation of tumor suppressor-like splicing programs. Together with other oncogenic alterations, these changes would help or drive the tumorigenic transition.

**Figure 5.**
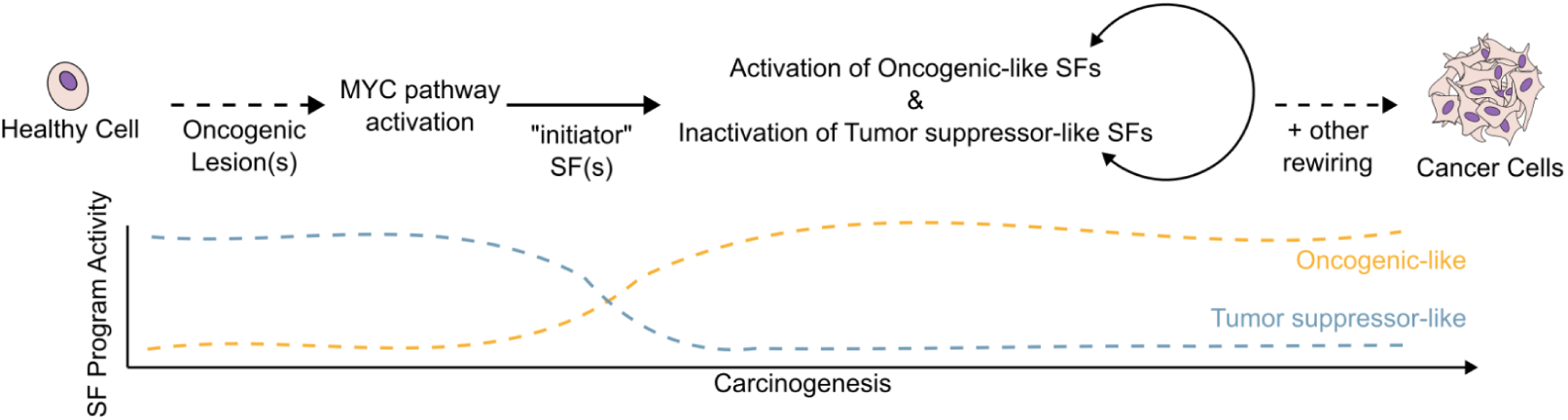
Model of coordinated regulation of cancer splicing programs during carcinogenesis.

This study adds to a growing body of evidence implicating MYC in the regulation of splicing factors in cancer^3,18,19,35,36^. While previous studies used bottom-up approaches to link MYC activation to splicing factor regulation, our systematic, top-down approach converged on the same conclusion, underscoring the robustness of our methodology and findings.

Our results further suggest that cancer splicing programs are not inherently cancer-specific but represent developmental cellular programs co-opted for tumor progression. However, their activation mechanisms may still be cancer-specific. For instance, MYC is frequently upregulated in cancers through copy number alterations^37^, whereas its developmental expression is tightly regulated by genome architecture and epigenetic mechanisms^38,39^. Additionally, carcinogenesis models demonstrated that both immortalizing and driver mutations are necessary to reprogram splicing activities and become tumorigenic. This underscores the critical role of cellular context in shaping carcinogenesis. Consequently, further research will be essential to delineate the prerequisites that set the stage for carcinogenesis.

Several aspects of our model will require further investigation. While MYC frequently drives cancer splicing programs, tumors without MYC activation may employ alternative pathways to regulate initiator splicing factors. Our data suggest that silencing diverse splicing factors can induce a switch in cancer splicing programs, with effects correlating with their proximity within PPI and splicing factor→exon networks (Fig. 2b). Thus, we foresee such cancers would also exhibit coordinated regulation of cancer splicing programs. Identifying these alternative regulators will be crucial for uncovering additional biomarkers and therapeutic targets. Furthermore, given the large number of target genes of master regulators like MYC, future studies should disentangle the relative contributions of splicing programs to the phenotypic changes of cancer cells compared to other MYC-driven molecular programs.

To conclude, our study establishes a robust framework for systematically investigating splicing factor activity regulation using high-throughput perturbation datasets. Applying this framework, we reveal that cross-regulatory interactions between splicing factors drive the coordinated activation of cancer splicing programs and that cancer cells co-opt MYC-regulated developmental splicing programs during carcinogenesis. We envision these findings pave the way for identifying novel genes and pathways regulating splicing programs, not only in carcinogenesis but across diverse biological processes.

## METHODS

### Quantification of splicing and gene expression from bulk RNA sequencing samples

All bulk RNA sequencing samples considered in this study were processed through the same pipeline. We downloaded raw sequencing reads and quantified gene expression and exon inclusion with *vast-tools*^41^. For each sample, we aligned sequencing reads to the hg38 genome assembly (Hs2 in VastDB) with the command “*vast-tools align --sp Hs2 --EEJ_counts --expr*”. We combined them using “*vast-tools combine --sp Hs2 --keep_raw_reads --keep_raw_incl --TPM*”. And, we set to NA the PSI of exons whose inclusion or exclusion was detected with less than 10 reads by running “*vast-tools tidy -min_N 1 -min_SD 0 --min_ALT_use 25 --noVLOW*”. For each dataset, this workflow results in a gene expression table corresponding to transcripts per million (TPM) –these values can range from 0 to potentially infinite– and an exon inclusion table quantifying the percentage of transcripts that include a certain exon or percentage spliced in (PSI) –these values can range from 0 to 100–. The details of each command can be found at https://github.com/vastgroup/vast-tools.

### Estimation of splicing factor activity using splicing factor networks and VIPER

VIPER is a computational method designed to infer protein activity based on molecular signatures^8^. VIPER uses regulatory networks that link molecular features (e.g., genes, exons) to their upstream regulators. By analyzing the differential regulation of these features between a reference and a perturbed condition, VIPER provides insights into the activity of regulators such as splicing factors, in our case. Specifically, VIPER tests for the enrichment of a regulator’s target within a rank-sorted molecular signature through rank-based enrichment analysis (aREA).

Estimation of splicing factor activity with VIPER requires two inputs: molecular signatures and regulatory networks. Molecular signatures represent the difference between conditions such as exon inclusion differences (e.g., deltaPSI) or gene expression differences (e.g., log2FC). Regulatory networks link each splicing factor to its targets, which can be exons for splicing factor→exon networks or genes for splicing factor→gene networks.

For all analyses, the *viper::viper* R function was executed with default parameters, ensuring consistent and reproducible results. VIPER processes the input molecular signature and regulatory network(s) to compute the activity scores for each splicing factor, which reflect their regulatory influence under the given conditions.

### Splicing factor→exon network inference

Splicing factor-exon networks were derived from two sources: data processed from Rogalska *et al.*^20^ and pre-existing networks available in the publicly accessible repository from Anglada-Girotto *et al.*^7^ (see Data Availability). Of note, Anglada-Girotto *et al.*^7^ analyzed a total of 64 different studies. These networks link splicing factors to their target exons in a functional manner as they are defined based on changes in exon inclusion observed upon splicing factor perturbation.

To generate splicing factor→exon networks from the perturbation experiments in Rogalska *et al.*^20^, we adopted the same pipeline described in Anglada-Girotto *et al.*^7^. Specifically, we quantified exon inclusion levels (PSI) for each sample using *vast-tools*, as previously described. PSI values were averaged across replicates for each experimental condition. For each splicing factor perturbation, we calculated the difference in PSI values (deltaPSI) between the perturbed and control conditions. Exon were considered targets of a perturbed splicing factor if their inclusion changed by at least 15 absolute units (|deltaPSI| ≥ 15). This threshold ensures that only substantial changes in exon inclusion are included in the network. The resulting splicing factor→exon interactions were formatted for use in VIPER, the tool used for splicing factor activity estimation from exon inclusion signatures. The sign of deltaPSI was used to define the mode of regulation (positive for activation, negative for repression), and the absolute value of deltPSI was used as the likelihood score for the interactions.

The repository provided by Anglada-Girotto *et al.*^7^ contains ready-to-use splicing factor→exon networks. These networks were generated following the same steps described above, ensuring consistency between the two data sources.

### Redefinition of cancer splicing programs

The inclusion of splicing factor→exon networks extended with data from Rogalska *et al.*^20^ enabled the estimation of splicing factor activities for 151 additional splicing factors compared to our previous study^7^. Thus, following the same approach, we redefined cancer splicing programs by identifying splicing factors that are recurrently activated or inactivated in primary tumor samples relative to their solid tissue normal counterparts.

We used exon inclusion matrices quantified with *vast-tools* from 14 TCGA cancer cohorts. Each cohort contained a minimum of 10 samples per condition (primary tumor and solid tissue normal). These data were obtained from Anglada-Girotto *et al.^7^*’s publicly available repository, which provides intermediate files (see Data Availability). Primary tumor exon inclusion signatures were computed by subtracting median exon inclusion values of solid tissue normal from those of primary tumor samples. Conversely, solid tissue normal exon inclusion signatures were computed by subtracting the median exon inclusion values of primary tumor samples from those of solid tissue normal samples. Using these exon inclusion signatures, we estimated splicing factor activities with VIPER, leveraging the splicing factor→exon networks.

We performed Mann-Whitney U tests using the Python function *scipy.stats.mannwhitneyu* to compare splicing factor activity between primary tumor and solid tissue normal samples within each cohort. An FDR-adjusted p-value threshold of 0.05 was used to identify significant differential activity.

We counted the number of cancer cohorts in which each splicing factor showed significant differential activity (either activation or inactivation). Splicing factors that were differentially active in more than five cohorts were categorized as oncogenic-like (recurrently activated), or tumor suppressor-like (recurrently inactivated).

### Cancer program activity

Splicing program-level activities were calculated as the median activity of all splicing factors associated with a specific program.

### Splicing factor→gene network inference

#### From bulk RNA sequencing data

Splicing factor→gene networks were constructed from bulk transcriptomic datasets using gene expression TPM matrices generated with *vast-tools*, as described above. These networks link splicing factors to their functional target genes –genes whose expression levels change upon perturbation of a specific splicing factor. Networks were derived both from raw data processed from Rogalska *et al.*^20^ and from pre-processed gene expression matrices available in the Anglada-Girotto *et al.*^7^ repository for intermediate files (see Data Availability).

Gene expression values in TPM were transformed into log2(TPM+1) to normalize and stabilize variance across genes with different expression levels. For each study, log-transformed gene expression values were averaged across replicates within the same experimental condition. Differential expression was calculated as the log2-fold change between the perturbed and control conditions for each splicing factor. Specifically, fold changes were determined by subtracting the average log2-transformed gene expression of the control condition from the corresponding perturbed condition. Genes were identified as targets of a given splicing factor if their expression levels changed by at least 1 absolute log2-fold change unit (|log2FC| ≥ 1) following splicing factor perturbation. The identified splicing factor→gene interactions were formatted for use in VIPER to estimate splicing factor activities from gene expression signatures. The absolute log2-fold change was used as the likelihood of the interaction, and the signal of the log2-fold change was assigned as the interaction’s mode of regulation (positive for activation, negative for repression).

#### From single-cell RNA sequencing data

Splicing factor→gene networks derived from single-cell transcriptomic data were constructed using three Perturb-seq experiments^10^: K562-essential, RPE1-essential, K562-genomic (see Data Availability). These experiments measured gene expression changes following the perturbation of thousands of genes, including different splicing factors (nK562-essential=248, nRPE1-essential=273, nK562-genomic=431). The networks link splicing factors to genes whose expression levels are altered upon splicing factor perturbation.

Raw read count matrices from single-cell data were aggregated into pseudo-bulk profiles by summing read counts of all cells labeled with the same perturbation. Pseudo-bulk counts were converted to counts per million (CPM) to account for differences in sequencing depth and log-normalized CPMs (log2(CPM+1)). For each dataset, we calculated log2-fold changes (log2FC) by comparing pseudo-bulk gene expression values under each perturbation condition to those in the control condition. Genes were included as targets in the splicing factor→gene networks if their expression changed by at least 1 absolute log2-fold change unit (|log2FC| ≥ 1) following perturbation. This threshold ensures that only genes with substantial expression changes are included. Finally, the splicing factor→gene networks were formatted for use in VIPER, setting absolute log2-fold changes as the likelihood of the interaction and corresponding signs as the mode of regulation of each interaction (positive for activation, negative for repression).

#### Combining bulk and single-cell networks

To create combined splicing factor→gene networks from bulk and single-cell RNA sequencing data, we collated all individual network files (generated separately from bulk and single-cell datasets) into a single directory. This directory was then used as input for VIPER. VIPER internally integrates all provided network files, combining their information to estimate a unified splicing factor activity.

### Making protein activity estimated with gene-based splicing factor networks equivalent to exon-based networks using shallow ANNs

#### Architectures of shallow artificial neural networks

To adjust splicing factor activities estimated from gene-based splicing factor networks to resemble those derived from exon-based networks, we developed two shallow ANN architectures using PyTorch:

- Element-wise scaling: this architecture applies a scaling factor to each splicing factor activity independently, essentially learning a unique adjustment for each factor.
- Fully connected layer: this architecture predicts the adjusted activity of each splicing factor by incorporating the estimated activities for all splicing factors, allowing for inter-factor dependencies in the adjustment.

#### Training the models

We used bulk RNA sequencing data from 1,019 cancer cell lines in the CCLE to train the models (see Data Availability). The training process involved the following steps:

1. *Data preprocessing*. Exon inclusion and gene expression were quantified using *vast-tools* as described above. The resulting matrices were median-centered by subtracting the median exon inclusion and gene expression values from each dataset to create normalized signatures.
2. *Splicing factor activity estimation.* Splicing factor activities were computed using VIPER with four types of networks, including exon-based and gene-based (from bulk, single-cell, or combined datasets) networks.
3. *Model training.* The models were trained for 20 epochs with a batch size of 64 and a learning rate of 0.01, using *nn.SmoothL1Loss* loss function. Input features were splicing factor activities derived from bulk, single-cell, or combined gene-based networks, while outputs were activities estimated using exon-based networks. We performed cross-validation with k=5, splitting the training dataset into five parts. In each iteration, four parts were used for training, and one part was left out for testing. This process was repeated five times, leaving out a different part each time to ensure robustness.

#### Evaluation

For each cross-validation split, type of gene-based activity, and ANN architecture, we computed Pearson correlation coefficients between the predicted splicing factor activities and the actual exon-based activities for both training and test splits. This evaluation assessed the models’ ability to generalize and reliably adjust gene-based splicing factor activities to match exon-based activities.

#### Inference across replicates of each model

Our training scheme produced five model replicates, each trained on a different data split from the cross-validation procedure. During inference for subsequent analyses, unless otherwise specified, splicing factor activities were adjusted using each model replicate independently. The outputs were then averaged to generate a consensus adjusted splicing factor activity matrix.

### Validation of carcinogenic regulation of cancer splicing programs

#### Bulk RNA sequencing dataset

We validated cancer splicing program activity switch detection using a bulk RNA sequencing dataset from Danielsson *et al.*, which measures the transcriptomic changes along the transformation of primary BJ fibroblasts into cancer cells through four distinct stages (untreated, immortalized, transformed, and metastatic)^21^ (see Data Availability). We processed raw data by quantifying exon inclusion (PSI) and gene expression (TPM) values using *vast-tools* as described above. Gene expression values were log-transformed for downstream analyses (log2(TPM+1)). Exon inclusion and gene expression signatures were computed by subtracting the average exon inclusion or gene expression profile of untreated fibroblasts from those of the other three transformation stages. Splicing factor activities were estimated using VIPER with various splicing factor→gene networks (bulk, single-cell, and combined) and splicing factor→exon networks. We also adjusted these activities using different pre-trained shallow ANNs with a fully-connected architecture. This analysis evaluated how well each network type recapitulated the coordinated activation of oncogenic-like splicing factors and inactivation of tumor-suppressor-like splicing factors, as observed with splicing factor→exon networks during carcinogenesis.

#### Single-cell RNA sequencing dataset

We also validated the detection of carcinogenic splicing switches using a single-cell RNA sequencing dataset from Hodis *et al.*, which studies the transformation of primary melanocytes into melanoma cells through the introduction of a series of mutations^24^ (see Data Availability). After downloading raw gene counts, we generated pseudo-bulk profiles by summing raw read counts across cells within each mutation condition. Gene expression profiles were normalized by computing CPMs and then log-transformed (log2(CPM+1)). We computed gene expression signatures by subtracting the gene expression profile of untreated melanocytes from those of each mutation condition. Finally, we estimated splicing factor activities using VIPER with bulk gene-based splicing factor networks, followed by adjustment with pre-trained fully connected ANN. Note that, for this dataset, reliable exon inclusion quantification was not feasible, so splicing factor→exon networks could not be applied.

### Splicing factor activity analysis of Perturb-seq data

To investigate how gene perturbations in the immortalized RPE1 cell line influence cancer splicing program activity, we analyzed preprocessed gene expression matrices from the Perturb-seq experiments^10^ previously used to generate single-cell splicing factor→gene networks. Splicing factor activities were estimated using log2-fold change gene expression signatures for each perturbation, derived from bulk gene-based splicing factor networks. These activities were then adjusted using the pre-trained fully connected ANN. Program-level activities were inferred as explained above, by computing the median splicing factor activity across oncogenic-like and tumor suppressor-like splicing factors.

### Pathway enrichment analysis of Perturb-seq data

To evaluate how each perturbation in the RPE1 Perturb-seq experiment affects pathway activities, we used the gene expression signatures as input for pathway enrichment analysis with the *clusterProfiler::GSEA* R function, using the MSigDB Hallmark gene sets. We used custom scripts for this analysis (see Code Availability), outputting NES values for each pathway and perturbation.

### Gene set enrichment analysis (GSEA) and overrepresentation analysis (ORA)

Unless stated otherwise, GSEA was performed using the *clusterProfiler::GSEA* R function and ORA using the *clusterProfiler::enricher* R function, both executed with default parameters. P-values were adjusted through the FDR method.

### Shortest path length analysis in STRINGDB PPI network

To analyze the distances between splicing factors in the STRINGDB PPI network, we downloaded the STRINGDB network (see Data Availability) and retained only interactions with a combined score above 900. Node names were standardized to gene symbols. For all splicing factors perturbed in the RPE1 Perturb-seq experiment, pairwise shortest path lengths were calculated using the *networkx.shortest_path_length* Python function.

### Identification of pathways regulating cancer splicing program activity switch during carcinogenesis

We first calculated Hallmark pathway NESs for both carcinogenesis models (Danielsson *et al.*, and Hodis *et al.*) using pre-computed gene expression signatures (see above). This analysis provided an enrichment score for each pathway under each condition along carcinogenesis.

To shortlist pathways mediating the connection between cancer driver alterations and the coordinated regulation of cancer splicing programs, we computed Pearson correlations between Hallmark pathway NESs and previously computed cancer splicing program activity differences across the two carcinogenesis models analyzed. This identified pathways that become active or inactive in parallel with cancer splicing program regulation. From these correlations, we shortlisted the top five positively and negatively correlated pathways.

We further refined the list by correlating the NESs of the shortlisted pathways with cancer splicing program activity differences in the RPE1 Perturb-seq dataset. This second prioritization aimed to differentiate true mediators from spurious correlations. Pathways mediating the splicing program switch are expected to be both regulated during carcinogenesis and responsive to gene perturbations in the Perturb-seq experiment, as perturbing these pathways would alter the transcriptome and, consequently, cancer splicing program activity. This approach ensures the identification of pathways relevant to both carcinogenesis and the regulation of cancer splicing programs.

### Analysis of Urbanski *et al.*^3^ dataset to validate MYC as a regulator of cancer splicing programs

Urbanski *et al.* analyzed the transcriptomic changes induced by activating MYC in the breast-derived MCF10A immortalized cell line. The system employed inducible MYC activation via 4-hydroxy tamoxifen (4-OHT) and performed bulk RNA sequencing at different time points (0, 8, 24, and 48 hours). Since 4-OHT itself can alter the transcriptome, the study also included a negative control, where the same cell line without the MYC-inducible system was treated with 4-OHT.

To evaluate how MYC activation influences cancer splicing programs, we downloaded the raw data (see Data Availability) and quantified exon inclusion (PSI) and gene expression (TPM) using *vast-tools* as described above. Gene expression data were log-transformed (log2(TPM+1)). Exon inclusion data were used to estimate splicing factor activities, and gene expression data were analyzed to compute NESs for MYC-related pathways across experimental time points.

#### Splicing factor activity analysis

Exon inclusion signatures were computed by subtracting the averaged time point 0 exon inclusion profiles from all the time points for each condition (with or without the MYC-inducible system). We used VIPER with splicing factor→exon networks to estimate splicing factor activities from these exon inclusion signatures. After averaging activities across replicates of each time point and condition, to account for transcriptomic changes caused by 4-OHT, we normalized splicing factor activities for each MYC-induced time point by subtracting the activities of the corresponding negative control. Normalized activities were aggregated into program-level activities by computing the median activity of splicing factors within each splicing program (oncogenic-like and tumor suppressor-like) for each time point.

#### Pathway analysis

We computed gene expression signatures by subtracting time point 0 expression values from all time points for each condition. These log2-fold change signatures were used to calculate NESs for MSigDB Hallmark pathways through GSEA. NES values for the MYC-induced condition were averaged across replicates and normalized by subtracting NES values from the corresponding negative control.

### Analysis of Cardoso-Moreira *et al.*^30^ dataset to assess the regulation of cancer splicing programs during developmental tissue differentiation

Cardoso Moreira *et al.* used bulk RNA sequencing to explore the transcriptomic changes in nine different tissues during developmental tissue differentiation. To analyze this dataset, we processed the raw sequencing reads to quantify exon inclusion (PSI) and gene expression(TPM) (see Data Availability). Gene expression values were log-normalized (log2(TPM+1)) for downstream analysis.

#### Splicing factor activity analysis

To perform splicing factor activity analysis, we computed exon inclusion signatures by subtracting the median exon inclusion across all samples within each tissue from individual sample values. We then estimated splicing factor activities using VIPER with splicing factor→exon networks. These activities were aggregated into cancer splicing program-level activities by computing the median activity across splicing factors within each program (oncogenic-like and tumor suppressor-like).

#### Pathway analysis

We computed gene expression signatures by subtracting the median gene expression across all samples within each tissue from the individual sample values. The resulting gene expression signatures were used to compute NESs for the MYC-related Hallmark pathways from MSigDB via GSEA.

## DATA AVAILABILITY

### Intermediate files generated from analyzing data analyses

We made available intermediate files generated throughout this study in this Figshare repository: https://figshare.com/articles/dataset/intermediate_files/28255892

### Previously published datasets used

*Consensus list of splicing factors from Anglada-Girotto et al.*^7^

Supplementary Table 1 in the publication.

*Splicing factor-exon networks from Anglada-Girotto et al.*^7^

Downloaded from https://github.com/MiqG/viper_splicing/tree/master/data/empirical_sf_networks-EX.

*Gene expression matrices from splicing factor perturbation experiments compiled in Anglada-Girotto et al.*^7^

Downloaded from https://doi.org/10.6084/m9.figshare.27835518.v1.

*Bulk RNA sequencing data of splicing factor perturbations from Rogalska et al.*^20^

We downloaded raw FASTQ files from the ENA website (PRJEB49033).

*Perturb-seq raw read counts from single-cell RNA sequencing in Replogle et al.*^10^

We downloaded raw read count matrices for three Perturb-seq screens targeting essential genes in K562 (“ReplogleWeissman2022_K562_essential.h5ad”) and RPE1 (“ReplogleWeissman2022_rpe1.h5ad”), and a genome-wide screen targeting all expressed genes in K562 (“ReplogleWeissman2022_K562_gwps.h5ad”) homogeneously processed by the scPerturb project^12^ (https://zenodo.org/records/7041849).

*The Cancer Genome Atlas (TCGA) exon inclusion matrices preprocessed in Anglada-Girotto et al.*^7^

Downloaded from https://doi.org/10.6084/m9.figshare.27835518.v1.

*Bulk RNA sequencing data of carcinogenesis from Danielsson et al.*^21^

We downloaded raw FASTQ files from the ENA website (PRJNA193487).

*Molecular data from the Cancer Cell Line Encyclopedia (CCLE) preprocessed in Anglada-Girotto et al.*^7^

Downloaded from https://doi.org/10.6084/m9.figshare.27835518.v1.

*Single-cell RNA sequencing data of carcinogenesis from Hodis et al.*^21^

We downloaded log-normalized reads (https://singlecell.broadinstitute.org/single_cell/data/public/SCP1334/engineered-melanocytes?filename=invitro_eng_melanoc_logTP10K.txt.gz) and corresponding metadata (https://singlecell.broadinstitute.org/single_cell/data/public/SCP1334/engineered-melanocytes?filename=invitro_invivo_all_metadatafile_mod_withCelltypes.csv) from their *in vitro* experiments transforming melanocytes into melanoma through mutagenesis from their website.

*List of mutational cancer driver genes (COSMIC-CGC)*

We downloaded the list of cancer driver genes from the COSMIC Cancer Gene Census website. https://cancer.sanger.ac.uk/cosmic

*STRINGDB protein-protein interaction network*

We downloaded the network from https://stringdb-static.org/download/protein.links.full.v11.5/9606.protein.links.full.v11.5.txt.gz. And, corresponding gene name aliases from https://stringdb-static.org/download/protein.aliases.v11.5/9606.protein.aliases.v11.5.txt.gz.

*CHEA Targets database*

We downloaded gene sets from the Harmonizome website: https://maayanlab.cloud/static/hdfs/harmonizome/data/chea22/gene_set_library_crisp.gmt.gz.

*MSigDB database*

We downloaded GO BP (https://www.gsea-msigdb.org/gsea/msigdb/download_file.jsp?filePath=/msigdb/release/2024.1.Hs/c5.go.bp.v2024.1.Hs.symbols.gmt), Reactome (https://www.gsea-msigdb.org/gsea/msigdb/download_file.jsp?filePath=/msigdb/release/2024.1.Hs/c2.cp.reactome.v2024.1.Hs.symbols.gmt), and Hallmarks (https://www.gsea-msigdb.org/gsea/msigdb/download_file.jsp?filePath=/msigdb/release/2024.1.Hs/h.all.v2024.1.Hs.symbols.gmt) gene sets from the MSigDB database.

*Bulk RNA sequencing data from Urbanski et al.*^3^

We downloaded raw FASTQ files from the ENA website (PRJNA754112).

*Bulk RNA sequencing data from Cardoso-Moreira et al.*^30^ *preprocessed in Anglada-Girotto et al.*^7^

Downloaded from https://doi.org/10.6084/m9.figshare.27835518.v1.

## CODE AVAILABILITY

The full data analysis pipeline of this manuscript is publicly available at https://github.com/MiqG/publication_regulation_cancer_asprograms and the corresponding standalone scripts to run VIPER with the splicing factor→exon and splicing factor→gene networks defined in this study at https://github.com/MiqG/splicing_factor_activity_analysis.

## ACKNOWLEDGEMENTS

We thank Sophie Bonnal, Jorge Herrero-Vicente, Jorge Ferrer, and the members of the Serrano and Valcárcel labs for the fruitful discussions and feedback provided throughout the development of the project. We also acknowledge the support of the Spanish Ministry of Science and Innovation through the Centro de Excelencia Severo Ochoa (CEX2020-001049-S, MCIN/AEI /10.13039/501100011033), the Generalitat de Catalunya through the CERCA programme and to the EMBL partnership. We are grateful to the CRG Core Technologies Programme for their support and assistance.

## AUTHOR CONTRIBUTIONS

Conceptualization: MA-G, SM-V, LS; Methodology: MA-G; Software: MA-G; Validation: MA-G; Formal analysis: MA-G; Investigation: MA-G; Resources: LS; Data Curation: MA-G; Writing - original draft: MA-G; Writing - review & editing: MA-G, SM-V, LS; Visualization: MA-G; Supervision: SM-V, LS; Project administration: MA-G; Funding acquisition: LS. All authors read and approved the final manuscript.

## SUPPLEMENTARY TABLES

**Supplementary Table 1. Redefined cancer splicing programs.**

